# RNA-seq gene profiling - a systematic empirical comparison

**DOI:** 10.1101/005207

**Authors:** Nuno A. Fonseca, John Marioni, Alvis Brazma

## Abstract

Accurately quantifying gene expression levels is a key goal of experiments using RNA-sequencing to assay the transcriptome. This typically requires aligning the short reads generated to the genome or transcriptome before quantifying expression of pre-defined sets of genes. Differences in the alignment/quantification tools can have a major effect upon the expression levels found with important consequences for biological interpretation. Here we address two main issues: do different analysis pipelines affect the gene expression levels inferred from RNA-seq data? And, how close are the expression levels inferred to the “true” expression levels?

We evaluate fifty gene profiling pipelines in experimental and simulated data sets with different characteristics (e.g, read length and sequencing depth). In the absence of knowledge of the ‘ground truth’ in real RNAseq data sets, we used simulated data to assess the differences between the “true” expression and those reconstructed by the analysis pipelines. Even though this approach does not take into account all known biases present in RNAseq data, it still allows to estimate the accuracy of the gene expression values inferred by different analysis pipelines. The results show that i) overall there is a high correlation between the expression levels inferred by the best pipelines and the true quantification values; ii) the error in the estimated gene expression values can vary considerably across genes; and iii) a small set of genes have expression estimates with consistently high error (across data sets and methods). Finally, although the mapping software is important, the quantification method makes a greater difference to the results.

## Introduction

Over the past five years, RNA-sequencing (RNA-seq) has been gradually overtaking microarrays to become the tool of choice for genome-wide analysis of the transcriptome [1]. Although RNA-seq offers multiple different perspectives on transcriptomic diversity (e.g., alternative splicing, allele-specific expression) one of the most important uses has been the quantification of gene expression levels or comparing them across conditions. However, processing the millions of short reads generated in a typical RNA-seq experiment to obtain measurements of gene expression levels requires considerable bioinformatics skills and computational resources. This process can be divided into three main steps (see Figure 1):

1. Filtering out reads that fall below some quality threshold
2. Aligning the reads to the genome or transcriptome
3. Quantifying gene expression levels

**Figure 1.**
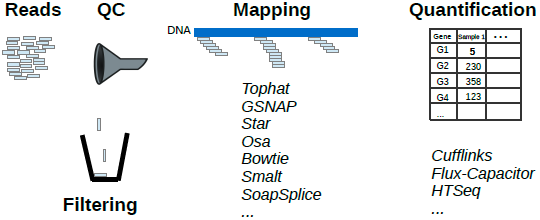
Gene profiling: from reads to gene expression.

The first step involves the identification of low quality reads (e.g., reads with biased base composition) that can be filtered out prior to mapping, thus avoiding potential mismatches and downstream errors. To perform the alignment, the user typically applies one of the many high-throughput sequencing mapping tools available [2], which requires choosing whether to align reads to the transcriptome or to the genome. Whilst the former is less computationally expensive, it is limited by the large number of reads that map to multiple transcripts of the same gene and by its reliance upon previously assembled annotation files. By contrast, mapping reads to the genome requires no knowledge of the set of transcribed regions or the way in which exons are spliced together. Given this greater flexibility, most approaches have mapped against the entire genome (where available) and we focus on this approach henceforth. In the third step, the aligned reads are passed to a quantification method [3–7] to obtain a measure of each gene’s expression (e.g., raw counts or normalized counts such as RPKMs [8]). Alternatively, tools for quantifying the expression of each transcript of a gene [7, 9–12] can be used and aggregated to obtain overall expression values.

Clearly, at each step of this process, the practitioner must select one of many possible options. A specific analysis pipeline is often selected based on subjective factors, such as popularity, ease of use, or past experience with some of the methods (possibly in a different context). It is unclear whether this choice, which is fundamental in many RNA-seq experiments, affects the measures of gene expression and hence downstream interpretation.

Previous studies have essentially compared set of alignment methods, typically used one quantification method, and have focused their results on assessing whether these approaches result in differences in the sets of differentially expressed genes identified. For example [13] assessed the ability to detect differentially expressed genes using three different aligners, a single quantification method (HT-Seq), and five differential expression methods by comparing the results to the ones obtained with microarrays. Other studies have assessed how different methods for identifying differentially expressed genes [13–15] perform when applied to the same input data sets. More recently, the RGASP consortium presented two broad evaluations of RNA-seq data analysis methods: one on the performance of several spliced alignment programs by focusing on the quality of the alignments [16]; the second study evaluated computational methods for transcript reconstruction and quantification from RNA-seq data [17]. In this work we address complementary yet fundamental questions: do different analysis pipelines affect the gene expression levels inferred from RNA-seq data? And, how close are the expression levels inferred to the “true” expression levels?

To address these questions we applied fifty RNA-sequencing pipelines (utilising all combinations of ten alignment methods and five quantification methods) to both real and simulated data and investigated whether this affected the estimates of gene expression levels. The alignment methods considered include both splice-aware (e.g., TopHat and Star) and splice-unaware (e.g., Bowtie) aligners in order to assess the effect of splice-aware aligners upon the inferred expression estimates. The quantification methods used were different versions or modes of HT-Seq, Cufflinks and Flux-capacitor.

A limitation of using real data to compare different pipelines for processing RNA-sequencing data is that no ground truth is known, and thus it is impossible to state that one method is more accurate than another. To address this limitation forty data sets were generated *in silico* with different characteristics (read length, coverage, single- and paired-end reads) to assess the error of each pipeline (how different is the output of each pipeline compared to the true expression values), the effect of sequencing depth on the results and whether longer reads lead to reduced errors.

## Results

### Pipeline choice affects measurements for real data

We first investigated how the choice of processing pipeline affected gene expression measurement levels obtained from four independent RNA-sequencing experiments (See Methods). For each of the 50 pipelines we obtained measures of gene expression levels and compared them across pipelines. We observed that, in general, the measurements of gene expression obtained from different pipelines were highly correlated, with a median Spearman correlation of 0.99 (Figure 2A and Supplementary Figure 1-3). Nevertheless, between any two pairs of pipelines, we observed that many genes had significantly different expression estimates (False Discovery Rate of 1%) with some genes having up to 10-fold differences in expression (for a specific comparison see Figure 2, panels C and D). This suggests that while the overall correlation in gene expression levels is relatively high, for a subset of genes different pipelines yield very different results.

**Figure 2.**
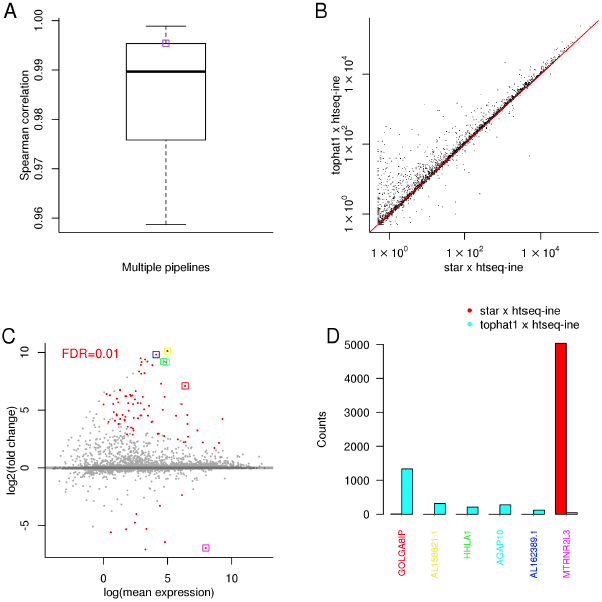
Illumina body map (E-MTAB-513) - RNA-seq data from Human. A) Spearman correlation distribution between the gene expression profiles inferred by different pipelines; B) correlation between two specific pipelines (the respective Spearman correlation is shown in plot A as a purple box); C) fold change between the gene expression values inferred by the same two pipelines - dots in red denote genes where the expression values are significantly different between the two selected pipelines (for a false discovery rate of 0.01); D) read counts inferred by the two pipelines for the six selected (boxed) genes in plot C).

### Simulated data

A limitation of using “real” data is that no ground truth exists, hence we next explored this more rigorously by performing a simulation study. We generated *in silico* data sets with different read lengths (50, 100, 150, 200 bp), overall coverage (10X, 30X, 60X, 120X) and with both single- and paired-end reads (see Materials and Methods for details). Note that each data set contains eight libraries. These data allowed us to assess how near the output of each pipeline was to the true expression values, the effect of sequencing depth on the results and whether longer reads lead to reduced errors. Obviously an ideal simulated data set should have properties similar to those observed in real data sets. However, to be able to compare all genes across different data sets thirty two data sets were generated *in silico* where all transcripts had approximately the same level of expression (referred as STE). In this setting the genes in a data set still have different raw expression levels that result from the stochastic nature of the simulator, different transcript lengths and number of transcripts, and sequencing depth. However, whilst having distinct advantages, these data sets (STE) do not recapitulate the expression patterns tipically observed in real data where a subset of all genes is expressed at varying levels. To address this potential limitation and validate any findings found using the STE data sets we generated *in silico* eight data sets (designated as MTE) in a way that resembles the expression patterns often observed in real data.

Overall, 82% of all pipeline runs (from a total of 1600) to analyze the STE data sets succeeded (see Figure 6 in Supplementary material for a breakdown by pipeline, read length, and depth). Some pipelines failed (during the mapping or quantification stage) therefore we only ranked the pipelines where more than 50% of runs succeeded. Using this criterion, four pipelines where Bowtie1 was used as the mapper were excluded from further analysis.

### Overall gene profiling comparison

To assess the various processing pipelines we used two different metrics: the Spearman correlation between the inferred and the true expression values; and the relative error (see Methods for more details). Figure 3 summarizes the results. We observed that, for all pipelines and data sets, the Spearman correlation between the true and inferred expression values is typically high (median of 0.92). The median error observed across all data sets and pipelines is slightly below 20%. When examining the sign of the error, i.e., whether the relative expression of a gene is over-or-under estimated when compared to the true expression values, we found that two third of genes had the gene expression levels underestimated while about one third overestimated (see Figure 10 in Supplementary material).

**Figure 3.**
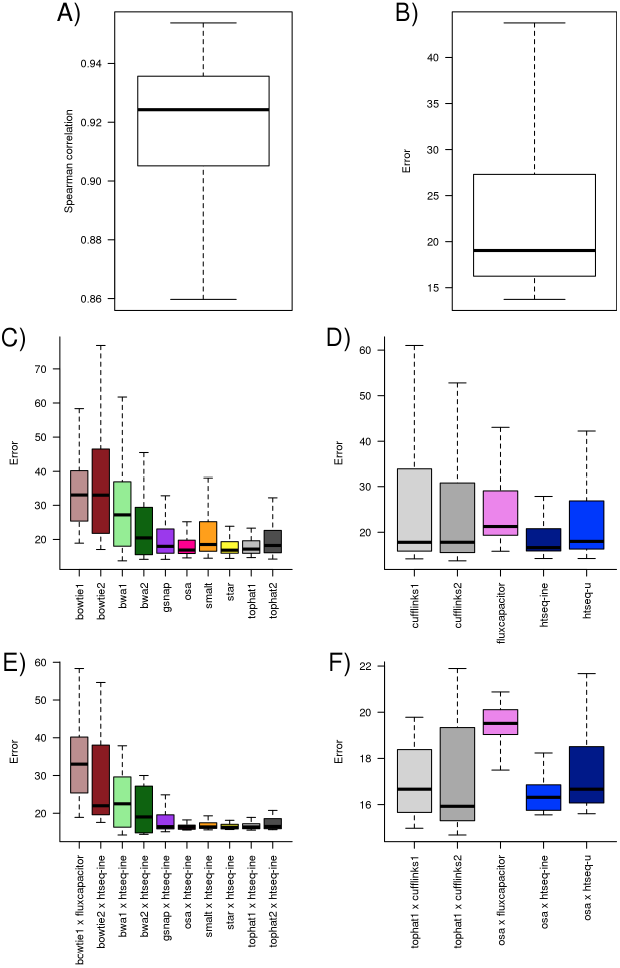
Distribution of Spearman correlation (A) and error (B) observed across all 32 data sets and all pipelines. Each observation corresponds to the value computed using a pipeline on a specific data set. The distribution of the error across all data sets and pipelines segmented by aligner (C) and quantification method (D). The distribution of the error across all data sets for the pipeline with lowest error for each aligner (E) and quantification method (F).

When we segment the error by the quantification method (Figure 3C) we observe that the median error for most quantification methods is close to the overall median error but the spread of the error varies largely. This suggests that some quantification methods (such as HT-Seq) are less dependent on the choice of the mapper than others.

To obtain an overall summary (Table 1), we ranked the pipelines for each data set (e.g., for the Spearman correlation we ranked the pipelines for each data set from 1 to 50 - from highest to lowest correlation). In the table it is shown the average rank for each pipeline, which was obtained by averaging the rankings of the pipeline across the 32 STE data sets. Therefore, since it is an average, it may happen that several pipelines have the same ranking or that no pipeline has an average rank of 1. An overall measure of performance, per pipeline, was obtained by summing the two individual rankings.

**Table 1.** Average rankings of the pipelines observed on the 32 data sets using two metrics (relative error and Spearman correlation). The overall rank was obtained by summing the average rank (N = 32) obtained on each metric. The average value and standard deviation across data sets is shown for each metric. The table is sorted by overall rank.

| Aligner | Pipeline | Overall Rank | Error |  | Spearman |  |
| --- | --- | --- | --- | --- | --- | --- |
| | Quant. Method | | Avg. Rank | mean ± sd | Avg. Rank | mean ± sd |
| osa | htseq-ine | 17 | 9 | 16.88 ± 2.00 | 8 | 0.94 ± 0.01 |
| tophat1 | htseq-ine | 18 | 10 | 17.28 ± 2.52 | 8 | 0.94 ± 0.01 |
| gsnap | htseq-ine | 23 | 11 | 19.87 ± 7.78 | 12 | 0.93 ± 0.01 |
| tophat2 | htseq-ine | 23 | 13 | 19.10 ± 5.93 | 10 | 0.94 ± 0.01 |
| osa | fluxcapacitor | 24 | 22 | 19.95 ± 2.04 | 2 | 0.95 ± 0.00 |
| star | htseq-ine | 24 | 11 | 17.37 ± 2.81 | 13 | 0.93 ± 0.01 |
| smalt | htseq-ine | 25 | 14 | 18.93 ± 7.22 | 11 | 0.93 ± 0.00 |
| tophat1 | fluxcapacitor | 27 | 23 | 19.79 ± 2.08 | 4 | 0.94 ± 0.00 |
| bwa2 | htseq-ine | 28 | 16 | 20.34 ± 6.05 | 12 | 0.93 ± 0.02 |
| star | fluxcapacitor | 29 | 23 | 20.51 ± 2.72 | 6 | 0.94 ± 0.00 |
| tophat1 | cufflinks2 | 31 | 13 | 22.74 ± 16.44 | 18 | 0.92 ± 0.03 |
| osa | cufflinks2 | 33 | 13 | 20.87 ± 9.57 | 20 | 0.92 ± 0.03 |
| star | cufflinks2 | 34 | 12 | 16.94 ± 2.96 | 22 | 0.92 ± 0.01 |
| tophat2 | fluxcapacitor | 34 | 26 | 22.17 ± 4.44 | 9 | 0.94 ± 0.01 |
| bwa2 | fluxcapacitor | 36 | 26 | 20.84 ± 2.99 | 10 | 0.94 ± 0.01 |
| osa | htseq-u | 36 | 16 | 22.07 ± 15.13 | 21 | 0.92 ± 0.02 |
| tophat1 | cufflinks1 | 36 | 13 | 22.35 ± 15.58 | 23 | 0.92 ± 0.01 |
| tophat1 | htseq-u | 36 | 17 | 19.84 ± 4.83 | 19 | 0.92 ± 0.00 |
| bwa2 | htseq-u | 37 | 20 | 23.75 ± 13.71 | 17 | 0.92 ± 0.02 |
| gsnap | cufflinks2 | 37 | 16 | 24.26 ± 17.44 | 22 | 0.91 ± 0.05 |
| gsnap | fluxcapacitor | 37 | 26 | 22.71 ± 6.43 | 11 | 0.94 ± 0.00 |
| smalt | htseq-u | 37 | 19 | 20.30 ± 7.67 | 19 | 0.92 ± 0.01 |
| osa | cufflinks1 | 38 | 13 | 21.12 ± 12.17 | 25 | 0.92 ± 0.01 |
| bwa1 | htseq-ine | 39 | 21 | 23.84 ± 8.04 | 18 | 0.92 ± 0.03 |
| star | cufflinks1 | 39 | 12 | 17.65 ± 4.28 | 28 | 0.91 ± 0.01 |
| tophat2 | cufflinks2 | 39 | 14 | 23.35 ± 16.97 | 25 | 0.91 ± 0.07 |
| gsnap | htseq-u | 40 | 16 | 23.46 ± 13.44 | 24 | 0.92 ± 0.01 |
| tophat2 | htseq-u | 40 | 19 | 21.44 ± 7.53 | 21 | 0.92 ± 0.01 |
| bwa2 | cufflinks2 | 42 | 22 | 25.93 ± 15.93 | 19 | 0.91 ± 0.05 |
| star | htseq-u | 42 | 18 | 18.48 ± 4.69 | 24 | 0.92 ± 0.00 |
| gsnap | cufflinks1 | 44 | 17 | 25.40 ± 17.08 | 27 | 0.91 ± 0.04 |
| bwa1 | htseq-u | 48 | 26 | 26.14 ± 8.57 | 22 | 0.91 ± 0.03 |
| tophat2 | cufflinks1 | 48 | 18 | 27.41 ± 21.56 | 30 | 0.90 ± 0.07 |
| smalt | cufflinks2 | 50 | 24 | 25.23 ± 15.73 | 26 | 0.90 ± 0.04 |
| bwa2 | cufflinks1 | 52 | 26 | 33.95 ± 22.11 | 27 | 0.88 ± 0.08 |
| smalt | cufflinks1 | 52 | 25 | 26.74 ± 16.89 | 28 | 0.90 ± 0.06 |
| smalt | fluxcapacitor | 52 | 28 | 26.59 ± 8.91 | 24 | 0.91 ± 0.03 |
| bwa1 | cufflinks2 | 53 | 27 | 31.39 ± 17.84 | 26 | 0.88 ± 0.07 |
| bwa1 | fluxcapacitor | 54 | 31 | 30.97 ± 11.13 | 23 | 0.89 ± 0.05 |
| bwa1 | cufflinks1 | 55 | 27 | 36.30 ± 22.90 | 28 | 0.87 ± 0.08 |
| bowtie2 | htseq-ine | 64 | 29 | 29.67 ± 12.57 | 35 | 0.86 ± 0.03 |
| bowtie1 | fluxcapacitor | 66 | 34 | 34.20 ± 11.43 | 32 | 0.87 ± 0.04 |
| bowtie2 | htseq-u | 68 | 32 | 32.96 ± 15.04 | 37 | 0.85 ± 0.03 |
| bowtie2 | fluxcapacitor | 72 | 35 | 34.49 ± 8.45 | 37 | 0.83 ± 0.05 |
| bowtie2 | cufflinks2 | 76 | 37 | 43.69 ± 21.38 | 39 | 0.82 ± 0.07 |
| bowtie2 | cufflinks1 | 77 | 36 | 40.13 ± 18.6 | 40 | 0.81 ± 0.05 |

Importantly, the error observed is independent of gene length. When the error is compared to the coverage of a gene the regression line of the error remains at the same level for different gene lengths (see Figure 7 in Supplementary material). This is consistent across different data sets and the same trend can be observed in multiple pipelines.

### Effect of aligners and quantification methods on gene profiling

When we segment the error by mapper (Figure 3D) it is unsuprisingly clear that error is higher for unspliced aligners (Bowtie 1 & 2, BWA 1 & 2). This difference is more pronounced in Supplementary Figure 4 where we observe that the error for unspliced aligners increased with read length. From the set of spliced aligners, Tophat1, OSA and Star show a very similar low median error rate and spread. These results suggest that these three mappers produce alignments with approximately the same quality. A similar observation was made in a previous study [25] that compared several methods for read alignment including TopHat and OSA.

Despite this, we observe that the overall pipeline rankings grouped the pipelines by quantification method, thus suggesting that the selection of the quantification method is more relevant than the (spliced) aligner used. It is also striking that one of the simplest quantification methods, HTSeq, when used in conjunction with different aligners shows up frequently towards the top of the rankings. Note that HTSeq discards all multi-mapping reads (i.e., reads that map to multiple locations) while Cufflinks and Flux-Capacitor attempt to allocate these reads to the most “probable” transcript/gene.

### Effect of sequencing data characteristics on gene profiling

Figure 5 shows the error distribution across all data sets and pipelines for different sequencing depths and segmented by read length. The median error decreases significantly (for Mann-Whitney-Wilcoxon test and a significance level of 0.05) when increasing depth from 10x to 30x. For larger depths the median error decreases but not significantly. Interestingly, if we restrict the analysis to data sets with read lengths of 50 or 100 bp there is no significant difference in the median errors when increasing the depth.

**Figure 4.**
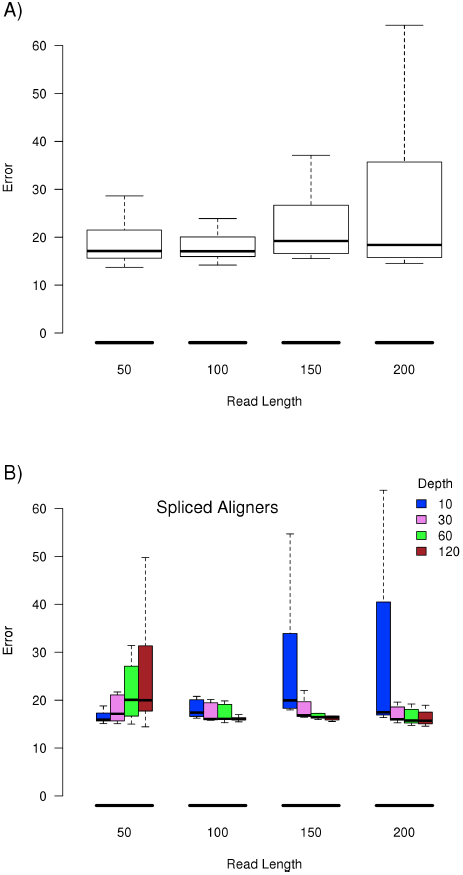
Distribution of the error across all data sets and pipelines for different read lengths (A), error observed using pipelines with spliced aligner on all data sets segmented by sequencing depth (B).

**Figure 5.**
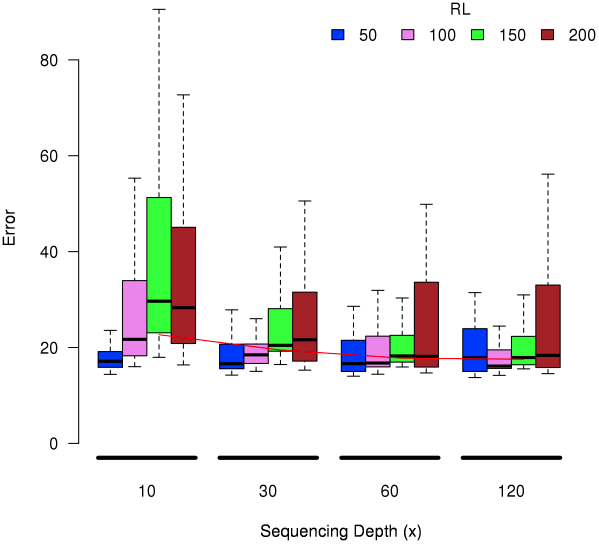
Distribution of the error across all data sets and pipelines for different sequencing depths and segmented by read length (RL).

**Figure 6.**
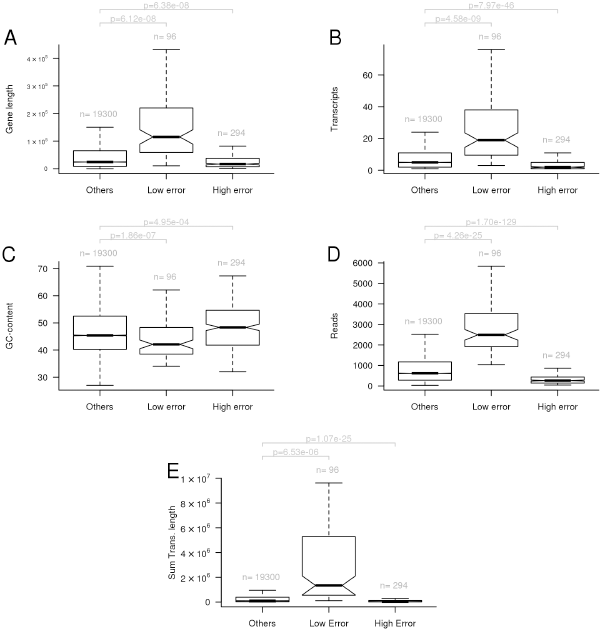
Genes with very high error (*>* 100%) and very low error (*<* 10%)]Genes with very high error (*>* 100%) and very low error (*<* 10%) across all data sets and pipelines combining OSA or Tophat1 with htseq-ine, Cufflinks2, and Flux-capacitor. For each group of genes it is shown the distribution of: A) gene length; B) number of transcripts; C) GC-content; D) true quantification (number of reads); E) sum of the length of transcripts of a gene. The p (p-value) was obtained by performing an unpaired t-test.

The rankings of the pipelines evaluated using paired-end data (a.k.a. pair-end tags) or single-end data are similar, having a Spearman correlation of 0.88 (see Supplementary Table 5 and Table 6). By comparing the two tables it is possible to observe that most pipelines in the top ranked positions, when considering data sets with single-end reads, are also in the top rank positions when considering the data sets with paired-end reads. Overall, the mean error observed was slightly higher in the data sets with paired-end data.

### Gene level analysis

An inspection of the results at gene level shows that a small subset of genes have consistently low/high errors across pipelines and data sets (Supplementary Figure 5). Figure 6 compares the set of genes with low and high errors to the remaining genes and gene characteristics (length, number of transcripts, GC-content, true read counts, and the sum of the length of all transcripts) in an attempt to understand if the set of genes with consistent high/low error could be explained by some characteristic.

The set of genes with consistent low error tend to have a high number of read counts. It is also noticeable that the set of genes with low error have longer gene lengths and more transcripts. However, this results directly from the simulated data since the number of true read counts are proportional to the sum of the length of the transcripts of a gene (which can be observed in panel E). This was confirmed in the additional eight simulated MTE data sets where there was no observed overlap between the genes with low error and the set of genes with consistent low error.

## Discussion

Overall the measurements of gene expression obtained from different pipelines are often highly correlated in the experimental data sets. However, between any two pairs of pipelines, many genes had significantly different expression estimates, thus different pipelines yield very different results for a subset of genes.

When considering the simulated data sets, the read counts inferred by the pipelines are highly correlated with the true values (see Figure 3). Since the data are simulated, the difference between the inferred read counts and the true values can be explained by reads that: i) had poor sequencing quality; ii) the aligner was unable to align; iii) the alignment or quantification method wrongly assigned to a gene. If the unaccounted or wrongly counted reads affect all genes in an unbiased way, then the relative expression values of the genes will remain close to the true values, and therefore lowering the error.

Table 1 shows the top ranked pipelines and demonstrates that there is no clear best pipeline across the metrics considered. For instance, the pipeline with OSA x Flux-capacitor has an average ranking of 22nd using the error metric, while having one of the highest average Spearman correlation. Note that these conclusions are based on simulated data.

Previous studies mentioned the critical role of the aligner [13,16] but our results show that the rankings are more often determined by the quantification method instead of the aligner used. This does not mean that the aligner used is unimportant, instead it indicates that the quality of (spliced) aligners may have reached a point where it does not appear to make a big difference which one is used in the context of gene profiling analysis. The preponderance of HT-Seq in the top positions is rather remarkable when one considers that it discards all alignments involving reads that map to multiple locations, as opposed to methods that use more complex algorithms to deal with such reads (such as Cufflinks and Flux-Capacitor). Possibly the more complex quantification methods may still have room for improvement.

Overall, increasing the sequencing depth reduced the quantification errors of the pipelines (see Figure 5), but only up to a certain point. The median error decreases significantly (for Mann-Whitney-Wilcoxon test and a significance level of 0.05) when increasing depth from 10x to 30x. For deeper depths the median error decreases but not significantly. When considering the additional eight MTE data sets the median error only decreases significantly when the sequencing depth was increased from 30x to 60x. For greater depths (120x) the error is not reduced. Previous studies [18] suggested that a rather stable detection of protein coding genes is reached at sequencing depths closer to 30x.

Regarding the set of genes with consistently high error across multiple data sets and pipelines, none of the gene characteristics considered (e.g., gene length, number of transcripts, gc-content, read counts) was a *sine qua non* condition to discriminate them (Figure 6). We next considered the size of the genes’ exons. The hypothesis was that the genes with high error could have shorter exons. Supplementary Figure 8 shows the distribution of the percentage of the length of the genes with exons smaller than 200 or 500 nucleotides does not differ between the set of genes with high error and the remaining genes. We also tested other possible hypothesis that could explain the list of genes with consistent high error based on the gene sequences, such as similarity to other genes (e.g., paralogous), repetitive sequences or shared subsequences. This can be condensed to the question of knowing how unique a gene’s sequence is, i.e., how many regions in the genome are similar to the exons of a gene. Supplementary Figure 9 shows that more than 50% of the genes with very high error also have higher number of similar regions in the genome than the set of remaining genes (see Material and Methods for details). However, 50% of the genes in the low error list also have a higher number regions in the genome similar to the genes’s exons. Furthermore, many of these genes also have high gene expression inference errors in the additional eight data sets where the range of expression values and number of expressed genes is closer to the values observed in real data. The list of genes is provided in Supplementary Tables 7, 8, and 9. The consistent high error observed for some genes across data sets and pipelines could not be predicted or discriminated based solely on the set of characteristics considered. Further analysis are required to determine if the same pattern is observed in other organisms. Nevertheless it may still be useful to consider this information when analysing Human RNA-seq data since several of these genes have been associated to various diseases (e.g., Urod [19], SNX5 [20], CAV2 [21]).

The median error observed across all data sets and pipelines is around 20%. The error does not affect all genes in the same way. The trend observed is that the error tends to get smaller with higher expression values (read counts). However, this does not mean that lower expressed genes will always have high errors. In fact we observed genes (see Figure 5) across all data sets and pipelines with an error close to 0% with either low or high expression values. Furthermore, genes with very high error but with either low or high expression values were also found, suggesting that other factors besides the expression values may affect the accuracy of the gene profiling methods.

## Conclusions

All in all, we have evaluated and compared gene profiling computational pipelines as a whole, from the initial sequencing reads to gene expression values. Fifty gene profiling pipelines were evaluated on thirty two *in silico* simulated data sets. The main findings can be resumed as follows:

1. The overall correlation between the quantification values inferred by the different pipelines and true values is above 0.90 and the error is around 20%. The error does not affect all genes in the same way and can vary considerably. A set of genes was found to have consistent high error (across data sets and pipelines) suggesting that other factors besides the expression values may affect the accuracy of the gene profiling methods.
2. The rankings indicate that quantification method used by a pipeline makes a greater difference in the results than the (spliced) aligner selected.
3. The read length effect on the performance of the pipelines varies depending if a spliced or unspliced aligner is used. The difference between using spliced and unspliced aligners is important - the error increases with the length of the reads with unspliced aligners.
4. The sequencing depth has an effect upon the errors of the pipelines. The overall error tends to stabilize around 30x/60x, hence deeper sequencing depths (more than 60x) will not yield more accurate results in the context of gene expression quantification.

There are several limitations with our approach. First, part of the results have been obtained using synthetic data and it is unclear to what extent these results are applicable to other experimental data sets. Although this question can arguably be raised for any kind of data used, simulated or experimental, we attempted to minimize this issue by using a large number of data sets. Second, we employed a single method to generate RNA-Seq data *in silico* which may introduce some bias. Thirdly, since we simulated RNA-seq from a single species (Human) it is possible that the pipelines evaluated may behave differently on data from evolutionary distant species. Lastly, this study does include several mappers and quantification methods but it is certainly not exaustive since many more mappers and quantification methods exist. In spite of these limitations, the results and conclusions presented have practical implications for RNA-seq data analysis.

## Materials and Methods

### Data

We analyzed the Illumina Human Body Map RNA-seq set (ArrayExpress accession: E-MTAB-513; http://www.ebi.ac.uk/arrayexpress) and three other experimental data sets. More details are provided in Supplementary Table 3.

Synthetic RNA-seq reads were generated *in silico* using Flux-Simulator (version 1.2.1-20130219021747) [22]. The Flux-simulator aims to provide an *in silico* reproduction of the experimental pipelines for RNA-Seq by using models based on analyzed RNA-Seq experiments from different cell types, sample preparation protocols and sequencing platforms.

Each simulated data set represents two conditions containing four replicates, where all transcripts have approximately the same level of expression. To achieve a sequencing depth of *D* ^1^, and assuming a transcriptome size of 60MB, the approximate number of reads (*nreads*) for a data set was defined as 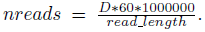 The following options were used in the generation of all data sets with Flux-simulator: FRAG_SUBSTRATE = RNA; FRAG_METHOD = UR; RTRANSCRIPTION = YES; RT_MOTIF = default; PCR_DISTRIBUTION = default; GC_MEAN = NaN; PCR_PROBABILITY = 0.05; FILTERING = NO; TSS_MEAN = 50; POLYA_SCALE = 100; POLYA_SHAPE = 1.5. FRAG UR ETA was set to 350 for reads smaller than 150 and 4x the read read length otherwise. By setting the GC_MEAN = NaN we disabled GC biases. The NB Molecules parameter was set to twice the number of reads. The Flux-simulator error models feature was used to simulate sequencing errors and base quality values. Instead of using the default error models available in Flux-simulator, we generated custom error models based on the reads from the experiment E-MTAB-513.

The Human genome (GRCh37.66) and respective annotation were obtained from Ensembl. The analysis focused on features annotated as protein coding genes and reads were simulated independently from all transcripts of each gene. Supplementary Table 3 summarizes the experimental data used and Supplementary Table 4 the synthetic data sets. The synthetic data set name includes information about the read length (length), sequencing depth (depth), and if the libraries are paired-end (pe) or single-end (se): *l*<*rl*>.*d*<*depth*>.<*se*|*pe*>. For instance, the data set with the name l100.d30.se contains single-end 100 base long reads with a sequencing depth of 30x.

An additional set of eight data sets was generated *in silico* using Flux-simulator as described above with the following differences: i) the range of expression values was automatically determined by Flux-simulator to mimic the range of expression values observed in experimental data; ii) the read lengths considered were 50 bp and 100 bp; iii) all libraries were single-end.

The data sets are publicly available at http://www.ebi.ac.uk/∼nf/gpeval/.

### RNA-seq gene expression pipelines

A RNA-seq pipeline for estimating gene expression is here considered as being composed by two steps: i) one tool for mapping the reads and generating the alignments; and ii) one quantification tool that takes the alignments and other information to infer the expression values of the genes.

There is a considerable number of tools for mapping high throughput sequencing reads and RNA-seq data in particular [2]. We included the following aligners in the evaluation: TopHat1 [23], TopHat2 [24], Osa [25], Star [26], GSNAP [27], Bowtie2 [28], Bowtie1 [29], Smalt ^2^, BWA1 [30], BWA2 [31]. Supplementary Table 2 summarizes which mappers are capable of aligning spliced reads (reads that span multiple exons).

The quantification tools considered include HT-Seq [3] (union mode/htseq-u and intersection non-empty mode/htseq-ine), Flux-capacitor [6], Cufflinks1 [4], Cufflinks2 [5].

The pipelines considered in the comparison include all combinations of selected aligners and quantification methods, which gives a total of 50 pipelines. All pipelines considered were implemented and executed using the iRAP RNA-seq pipeline [32]. Many of the methods that are compared in this paper allow the user to select the value of certain parameters, that can affect the results in various ways. We have mostly used the default values provided in the iRAP pipeline. Supplementary Table 1 summarizes the parameter values used for each tool. For information about the meaning of the different parameters, we refer to the manuals and original publications describing the respective methods.

### Methodology

The following metrics were used to compare the expression estimates inferred by different pipelines given the same data set:

- Spearman correlation between the true and inferred read counts per gene;
- Relative error.

The error metric measures how different the inferred quantification values are from the true values. The Spearman correlation allows an investigation of whether the ranking of gene expression levels in the inferred and true data sets are consistent. Arguably, the most appropriate metric to evaluate the accuracy of the pipelines is their *error*. For a given data set, let *G* be the number of genes, *S* the number of samples, *C* the number of conditions, *P* the number of pipelines, and *o*_*gscp*_ the number of reads assigned to gene *g* (1, …, *G*) in sample *s* (1, …, *S*) for condition *c* (1, 2) using pipeline *p* (where *p* ∈ 1, …, *P* denote the index of the pipeline). For the simulated data sets the true quantification value is known and it is denoted as *t*_*gsc*_. *N*_*ps*_ is the total number of reads of sample *s* assigned by the pipeline *p*.

The quantification error for a gene *g* using pipeline *p* is calculated as:

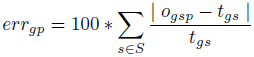

The overall quantification error of pipeline *p* is defined as

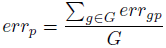

The formulation does not acknowledge that some reads may be discarded in some stage of the analysis pipeline and, therefore, not included in the quantification. To tackle this issue we propose the relative (normalized) quantification error for a gene *g* using a pipeline *p*, which is calculated in an analogous way:

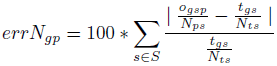

The relative quantification error is often referred to simply as error. In practice, to handle the cases where the true quantification of a gene is zero (reads), the error is computed by adding a 1 to the denominator and numerator.

The overall quantification error of a pipeline *p* is defined as the median error of all genes.

Finally, a note about the read counts for Cufflinks. Cufflinks does not output read count; instead it produces, amongst many values, the read coverage by transcript. Therefore, pseudo-counts were produced based on read coverage for each transcript and then aggregated to obtain the pseudo-counts per gene.

### Rankings

The pipelines were ranked using the two metrics mentioned above, separately for each data set. The average rank of a pipeline, using each of the two metrics, was computed as the mean rank of the pipeline across all data sets. The sum of the rankings of each metric was used to create an overall rank. The mean overall rank was determined when aggregating the results across data sets.

### Uniqueness of gene sequences

A gene’s sequence “uniqueness” is calculated based on how many regions in the genome are similar to the genes’s exons. The similarity of each exon was computed by aligning each exon to the genome using BLAST (version 2.2.12). Only high-scoring segment pairs (HSP) with an e-value below 0.01 and more that 90% identity, were considered. The alignments were later filtered by length (greater than 30 or 150 nucleotides). The uniqueness of a gene *g* with exons *e*_1_*, …* is computed as the *max*(*na*(*e*_1_)*, …*) where *na*(*e*) is the number of alignments obtained by BLAST for an exon.

## Acknowledgments

The authors would like to thank Konrad Rudolph, Robert Petryszack, Karyn Megy, and the members of the functional genomics team for their feedback.

## Footnotes

1 Transcriptome sequencing depth represents the average number of times a given base in the transcriptome is sequenced.

2 Smalt: http://www.sanger.ac.uk/resources/software/smalt/

